# Protein Language Model–Conditioned Graph Neural Networks for Multitask Ligand Activity Prediction Across Human GPCRs

**DOI:** 10.64898/2026.09.27.754816

**Authors:** Manashi De, Ekarsi Lodh, Shalini Majumder, Tapan Chowdhury

## Abstract

Predicting ligand activity across G protein-coupled receptors (GPCRs) requires models that capture both molecular structure and receptor-specific information while remaining robust to chemical and target-domain shift. We developed a multimodal graph neural network that combines explicit ligand molecular graphs with frozen protein language model representations of GPCR sequences and jointly predicts quantitative pActivity and binary activity. The model was trained on 271,739 curated ligand–GPCR pairs spanning 183,694 ligands and 216 human GPCRs and evaluated using random, Bemis–Murcko scaffold, and strict cold-ligand partitions. With ESM-2 650M receptor embeddings, the selected model achieved mean absolute errors of 0.513 ± 0.006, 0.540 ± 0.006, and 0.641 ± 0.005 pActivity units under random, cold-ligand, and scaffold evaluation, respectively, substantially outperforming a protein-aware fixed-feature multilayer perceptron and consistently improving quantitative prediction over a matched GINE reference. Removing receptor embeddings markedly degraded both regression and classification, whereas differences among ESM-2 35M, ESM-2 650M, and ProtT5 were comparatively small. Independent evaluation on 6,319 ChEMBL 37/BindingDB pairs revealed a substantial external domain shift, with mean absolute error increasing to approximately 0.94–0.95 despite chemically stringent internal validation. Nevertheless, receptor-specific information remained highly informative in a DRD2–DRD3 selectivity analysis: the cold-ligand ensemble achieved *R*^2^ = 0.853, Spearman *ρ* = 0.916, and ROC-AUC = 0.966 for strong DRD3 selectivity, whereas the receptor-independent ablation approached chance-level discrimination. These results establish protein-language-model-conditioned molecular graph learning as a scalable strategy for GPCR bioactivity prediction while identifying chemotype novelty, external-domain transfer, and richer ligand–receptor interaction representations as key remaining challenges.

## 1 Introduction

G protein-coupled receptors (GPCRs) constitute one of the largest and most pharmacologically important families of human membrane proteins. Approximately 800 GPCR genes are encoded in the human genome, where they translate chemically diverse extracellular signals into intracellular responses controlling neurotransmission, sensory perception, cardiovascular function, metabolism, immunity, endocrine regulation, and numerous other physiological processes [1], [2]. Their therapeutic importance is exceptional: a recent systematic analysis identified 516 approved drugs acting at 121 GPCRs, corresponding to approximately 36% of approved drugs, while a further 337 GPCR-directed agents spanning 133 receptors are under clinical investigation [1]. Yet only about one-third of non-sensory GPCRs are currently targeted by approved therapeutics, leaving a substantial fraction of the GPCRome pharmacologically underexploited.

The tractability of GPCRs as drug targets does not make ligand discovery straightforward. GPCRs are conformationally dynamic proteins whose pharmacology depends not only on ligand occupancy but also on receptor state, binding-site geometry, allosteric communication, signalling bias, and subtype-specific interactions [3]. Experimental structural biology has transformed the field, and resources such as GPCRdb now integrate hundreds of experimentally determined receptor structures with state-specific AlphaFold-derived models and extensive ligand annotations [2]. Nevertheless, structural coverage remains uneven across the receptor family, and a single static structure cannot fully represent the ligand-dependent conformational ensembles that govern GPCR activation and selectivity [2], [3]. These limitations are particularly relevant for poorly characterized or orphan receptors and motivate computational approaches that can exploit the much broader availability of protein sequence and bioactivity data without requiring an experimentally resolved ligand–receptor complex.

At the same time, public chemogenomic resources have reached a scale at which family-wide modelling has become increasingly feasible. ChEMBL contains standardized bioactivity measurements across large numbers of compounds and protein targets [4], [5], BindingDB provides an independently curated source of protein–small-molecule activity data [6], and UniProt supplies standardized receptor sequences and annotations [7]. GPCR-specific resources have similarly expanded: the 2023 GPCRdb release integrated more than 200,000 ligands and hundreds of thousands of potency or affinity measurements from ChEMBL, Guide to Pharmacology, PDSP *K*_*i*_, and related sources [2]. Such resources make it possible to move beyond target-by-target QSAR models toward architectures that learn transferable relationships across ligands and receptors. They also introduce substantial heterogeneity, however, because pharmacological measurements such as *K*_*i*_, *K*_*d*_, *IC*_50_, and *EC*_50_ reflect related but non-equivalent experimental quantities collected under different assay systems and conditions. The resulting modelling problem is therefore simultaneously one of representation learning, multitask supervision, and domain generalization.

Computational protein–ligand modelling has evolved rapidly in response to these challenges. Early data-driven affinity models relied primarily on engineered molecular and protein descriptors, similarity kernels, or conventional machine-learning algorithms. DeepDTA subsequently demonstrated that continuous drug–target affinity could be learned directly from one-dimensional ligand and protein representations using convolutional neural networks [8]. GraphDTA replaced the ligand string representation with an explicit molecular graph, showing that graph neural networks (GNNs) could capture molecular topology more naturally and improve affinity prediction relative to sequence-only ligand encodings [9]. More recent approaches span ligand and protein language models, molecular graphs, knowledge graphs, three-dimensional complexes, and hybrid sequence–structure architectures, reflecting the broader recognition that the central problem is not simply how to encode a molecule or a protein in isolation, but how to represent their interaction in a way that transfers beyond the combinations encountered during training [10].

Molecular graphs are particularly attractive for ligand representation because atoms and covalent bonds provide a direct relational description of chemical structure. Message-passing architectures can learn local chemical environments and propagate information across the molecular topology without relying entirely on predefined fingerprints. Their superiority is nevertheless not universal. Systematic comparisons have shown that conventional fingerprints remain competitive on many molecular-property tasks, especially in smaller data regimes, and that performance can deteriorate sharply when models are evaluated on previously unseen scaffolds [11]. The value of graph learning must therefore be established against strong fixed-feature baselines and under chemically meaningful split strategies rather than inferred from architecture alone. Protein representation has undergone a parallel transformation through self-supervised protein language models (PLMs). Models such as ESM-2 [12] and ProtT5 [13] are pretrained on millions of protein sequences and encode evolutionary, structural, and functional regularities without requiring task-specific structural annotations. GPCR-BERT has further shown that transformer-derived sequence representations can recover biologically meaningful organization within the GPCR superfamily itself [14]. In protein–ligand prediction, G-PLIP integrates chemical and protein information for structure-free bioactivity prediction [15], PSICHIC derives physicochemical ligand–protein interaction fingerprints directly from sequence information [16], and PMMR uses pretrained multimodal representations to improve transfer under cold-start drug–target binding scenarios [17]. Collectively, these studies establish PLMs as powerful target encoders, while leaving open how best to combine a global receptor representation with explicit ligand graph learning at family scale.

This question is especially timely for GPCRs. Recent GPCR-focused models have begun to move beyond single-receptor activity prediction toward broader receptor panels. AiGPro uses multi-scale sequence processing and bidirectional cross-attention to model agonist and antagonist bioactivity across 231 human GPCRs, demonstrating the feasibility of large-scale family-level potency prediction [18]. BOLD-GPCRs more recently combined transformer-derived receptor information with neural classifiers to predict ligand bioactivity and mutational effects across Class A GPCRs [19]. These methods represent important advances over earlier models restricted to individual targets or small GPCR subsets. Their objectives, representations, and evaluation paradigms nevertheless differ substantially: AiGPro is centred on agonist/antagonist functional activity, whereas BOLD-GPCRs emphasizes Class A bioactivity classification and mutation effects. This leaves complementary questions regarding whether explicit molecular-graph learning and PLM receptor representations can support a unified quantitative-and-binary activity model across a broad human GPCR panel, and how such a model behaves when chemical similarity or database provenance is deliberately shifted.

A second unresolved issue concerns the relationship between quantitative activity prediction and binary ligand classification. Historically, drug–target interaction models were frequently formulated as active/inactive classifiers, whereas affinity models such as DeepDTA and GraphDTA focused on continuous measurements [8], [9]. These formulations discard different types of information: thresholding continuous measurements removes quantitative ordering, whereas regression alone does not provide an explicit activity decision for screening. More recent approaches have begun to couple the two objectives. SLAB, for example, jointly learns binding-affinity regression and interaction classification [20], and Multi-task Bioassay Pre-training exploits related quantitative and ranking objectives across heterogeneous assay data [21]. For GPCR datasets, an additional complication arises from compounds whose activities lie close to an arbitrary active/inactive threshold. A multitask formulation that retains these measurements for regression while masking uncertain binary labels offers a natural way to preserve quantitative information without forcing borderline interactions into an artificial class.

The credibility of such models depends equally on how they are evaluated. Random pair-level splits can permit substantial chemical similarity between training and test compounds and may therefore overestimate performance expected for prospective chemistry. Scaffold-aware evaluation provides a more stringent estimate of transfer to distinct chemotypes, while strict ligand-disjoint splits test generalization to compounds never observed during training [11]. Even these grouped internal evaluations, however, need not reproduce the distribution shift encountered when models are applied to later database releases, independently curated measurements, or previously unseen targets. ZeroBind explicitly demonstrated the difficulty of unseen-drug and unseen-protein prediction [22], while PMMR similarly focused on improving cold-start transfer [17]. Independent temporal or source-separated testing therefore provides information that cannot be recovered from internal cross-validation alone. This distinction is particularly important in GPCR modelling, where activity measurements combine multiple pharmacological endpoints and where receptor coverage is highly uneven across the family.

Receptor subtype selectivity provides an additional test of whether a model has learned genuinely target-conditioned information rather than predominantly ligand-side correlations. Closely related GPCRs can share substantial sequence and binding-site similarity while differing sufficiently in local interactions to produce pharmacologically important selectivity profiles. DRD2 and DRD3 provide a representative example: subtype-selective dopamine-receptor ligands remain an active medicinal-chemistry problem, and recent studies have used machine learning, docking, and experimentally guided structure–activity analyses to identify determinants of D2/D3 preference [23], [24], [25], [26]. A family-wide model that predicts each receptor independently therefore offers a stringent test: correct selectivity must emerge from the difference between two receptor-conditioned predictions rather than from a receptor-pair-specific selectivity model.

To address these limitations, we developed a GPCR-focused multimodal graph-learning framework that combines explicit lig- and molecular graphs with frozen protein language model representations of receptor sequences and jointly predicts quantitative pActivity and binary interaction status. The framework was developed using a rigorously curated ChEMBL 36 dataset comprising 271,739 ligand–GPCR pairs, 183,694 unique ligands, and 216 human GPCRs, with *K*_*i*_, *K*_*d*_, *IC*_50_, and *EC*_50_ measurements harmonized on a common pActivity scale. Borderline interactions were retained for quantitative regression while being masked from binary classification, allowing both tasks to share a common representation without imposing uncertain activity labels. Ligands were encoded using a residual edge-conditioned graph neural network and fused with frozen ESM-2 or ProtT5 receptor representations, with a protein-aware fixed-feature MLP and a matched PyG-GINE implementation providing complementary baselines. Generalization was evaluated under random, Bemis–Murcko scaffold, and strict cold-ligand partitions across three independent training seeds, together with receptor-representation and single-task ablations. To assess transfer beyond the development distribution, the selected model was evaluated without retraining on an independently curated, leakage-audited ChEMBL 37/BindingDB test set and stratified by ligand, scaffold, target, and double-cold novelty. Receptor-specific information learned across the GPCR family was further examined through DRD2–DRD3 selectivity prediction, complemented by receptor ablation, fragment masking, and atom-level differential attribution.

The principal contributions of this work are as follows:

i. A GPCR-specific multitask learning framework that jointly models quantitative ligand activity and binary interaction status while integrating molecular graphs with pretrained receptor sequence representations.
ii. A controlled evaluation of model representation and architecture, separating the effects of molecular graph learning, receptor embeddings, graph architecture, and multitask optimization.
iii. A stringent generalization framework spanning random, scaffold-disjoint, cold-ligand, and independent external evaluation, including chemically and biologically defined novelty strata.
iv. A receptor-subtype validation of learned target information through DRD2–DRD3 selectivity prediction from a single GPCR-wide model.
v. Ligand-level interpretation of receptor-conditioned predictions using fragment perturbation and differential atom attribution.

The remainder of the paper is organized as follows. Section 2 describes GPCR and bioactivity curation, ligand graph construction, PLM receptor encoding, the CustomGNN and multitask objective, internal partitioning and controls, comparator models, external-set construction, and the DRD2–DRD3 analysis. Section 3 presents and discusses model selection, internal chemical-space generalization, the contributions of receptor representation and graph architecture, multitask effects, independent external performance and its error structure, and receptor-subtype selectivity and interpretation. Section 4 summarizes the principal conclusions and the implications for future GPCR bioactivity modelling.

## 2 Methods and Materials

### 2.1 Problem Formulation and Study Design

The study was formulated as a multimodal ligand–target prediction problem in which each observation corresponds to a small-molecule ligand (*L*_*i*_) paired with a G protein-coupled receptor (GPCR) target (*P* _*j*_). The ligand was represented as a molecular graph (*G*_*i*_ = (*V*_*i*_, *E*_*i*_)), whereas the receptor was represented from its amino acid sequence using a pretrained protein language model (PLM). Two related prediction tasks were learned jointly: continuous regression of ligand–GPCR activity and binary classification of active versus inactive interactions.

For ligand–receptor pair (*i, j*), the multitask observation was represented as:

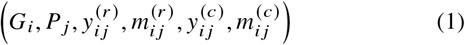

where 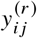 denotes the *continuous pActivity target*, 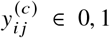 denotes the *binary activity label*, and 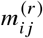 and 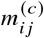 are task-specific masks identifying observations valid for regression and classification, respectively. This masking scheme allowed measurements in the intermediate activity range to contribute to quantitative regression without imposing an uncertain binary activity label.

Model development was performed using three complementary internal splitting schemes: random pair-level splitting, Bemis–Murcko scaffold splitting, and strict cold-ligand splitting. A multilayer perceptron (MLP) using fixed molecular features served as the principal baseline. In addition to the custom graph neural network (CustomGNN), a GINE-based PyTorch Geometric [27] implementation was evaluated as an architectural reference implementation rather than as the primary baseline. Protein representation variants and single-task ablations were evaluated during internal model development. External evaluation was subsequently performed for the GNN models using independently curated data; manuscript-level external performance is reported for the selected CustomGNN configuration. The biological utility of the selected model was further examined using a held-out *Dopamine receptor* D2 (DRD2) and *Dopamine receptor* D3 (DRD3) ligand-selectivity case study.

### 2.2 GPCR Target Identification and Bioactivity Data Acquisition

The internal dataset was constructed from ChEMBL release 36 [4], [5]. Human targets annotated as SINGLE PROTEIN were first extracted from the ChEMBL relational database together with target identifiers, target-component mappings, UniProt accessions [7], protein sequences, protein-class annotations, and target descriptions. GPCRs were identified from ChEMBL protein-classification information and associated target annotations using GPCR-related descriptors encompassing canonical G protein-coupled receptor, seven-transmembrane, rhodopsin-like, secretin-like, metabotropic glutamate, frizzled, adhesion, olfactory, and taste receptor annotations.

Quantitative ligand–target measurements were extracted for four pharmacologically relevant activity types:

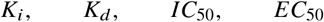

Only assays with a ChEMBL target-confidence score of at least 7 were retained. Activity records were required to contain a numerical standard value and standard unit. Canonical SMILES [28], [29] and standard InChIKey [30] identifiers were additionally required for downstream molecular standardization and pair construction.

Activity values were normalized to molar units. Supported units comprised molar, millimolar, micromolar, nanomolar, and picomolar measurements. For a quantitative activity value (*A*) expressed in molar units, pActivity was defined as:

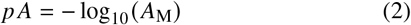

When a ChEMBL pChEMBL value was available, this standardized value was retained; otherwise, pActivity was calculated directly from the normalized molar activity using Eq. (2). Measurements with (*p A* < 2) or (*p A* > 12) were excluded. ChEMBL records flagged as potential duplicates were removed. For construction of the quantitative ligand–target dataset, inequality-censored activity relations were excluded; records represented by equality, approximate equality, or an unspecified equality relation were retained.

### 2.3 Ligand–GPCR Pair Aggregation and Multitask Target Construction

Activity measurements were aggregated at the level of the unique standardized ligand–GPCR pair, identified by the combination of standard InChIKey and UniProt accession. If (*n*_*ij*_) eligible measurements were available for ligand (*i*) and receptor (*j*), the regression target was defined as the median pActivity:

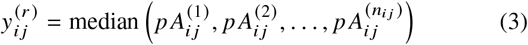

The mean, standard deviation, number of contributing activity records, contributing assay identifiers, and activity-type provenance were retained for quality control and downstream analysis. Median aggregation was chosen to reduce sensitivity to individual extreme experimental measurements while preserving a single ligand–target response value for supervised learning.

Binary labels were derived from the aggregated pActivity using an activity threshold of 6.0 and an inactivity threshold of 5.0:

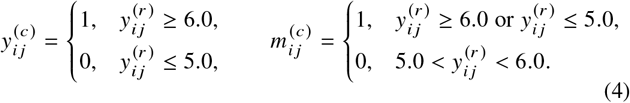

Pairs in the interval (5 < *p A* < 6) were therefore considered ambiguous for binary classification and assigned 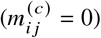, while remaining valid regression observations with 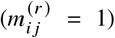. This design avoids forcing borderline ligand–receptor interactions into either binary class while retaining their quantitative information.

To limit extremely sparsely represented receptors, GPCRs with fewer than 50 curated ligand–target pairs were removed. The resulting internal multitask dataset contained 271,739 ligand–GPCR pairs involving 183,694 unique ligands and 216 GPCR targets. All 271,739 observations had valid regression targets. Of these, 225,527 were valid for binary classification, comprising 212,492 active interactions (94.2%) and 13,035 inactive interactions (5.8%); the remaining 46,212 observations were retained only for regression.

Table **??** summarizes the principal curation criteria and resulting multitask dataset composition, while Algorithm **??** provides the complete bioactivity-curation and multitask-label construction workflow.

#### Algorithm 1

GPCR bioactivity curation and multitask dataset construction

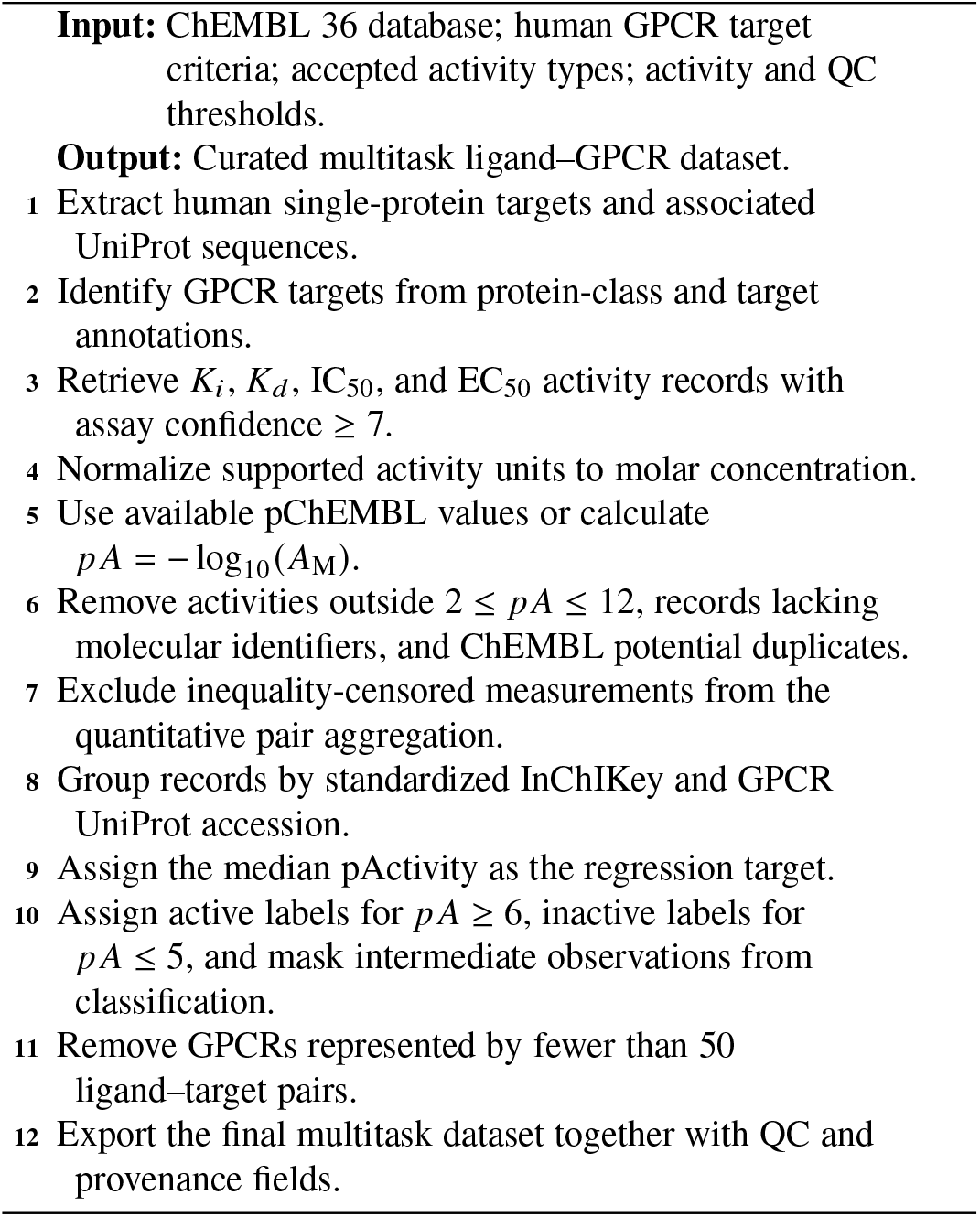

### 2.4 Ligand Molecular Representation

#### 2.4.1 Molecular Standardization

Unique ligands were identified using standard InChIKeys and canonical SMILES. Molecular structures were parsed with RDKit [31] and canonicalized using isomeric canonical SMILES. For multicomponent structures, the largest molecular fragment was retained during ligand feature preparation. Invalid structures were recorded separately and excluded from the graph/model index.

Two forms of ligand representation were generated: fixed cheminformatics features for the MLP baseline and explicit molecular graphs for the GNN models.

### 2.4.2 Fixed Features for the MLP Baseline

The fixed ligand representation contained a set of RDKit physico-chemical descriptors, along with a 2048-bit Morgan fingerprint of radius 2 (approximately equivalent to ECFP4) [32], both generated using RDKit. The descriptor panel included molecular weight, lipophilicity, topological polar surface area, hydrogen-bond, rotatable-bond, ring, heteroatom, charge, structural-complexity, and related physicochemical quantities. The complete descriptor definitions are provided in Supporting Information Table S1.

For protein-aware MLP experiments, the ligand descriptor and fingerprint vectors were concatenated with the corresponding GPCR PLM embedding. Feature standardization was fitted using the training partition only and then applied unchanged to validation and test observations. The regression target was similarly standardized using training-set statistics during MLP optimization and transformed back to the original pActivity scale for evaluation.

### 2.4.3 Molecular Graph Encoding

For the GNN models, each ligand was represented as:

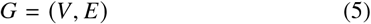

where each node (*v* ∈ *V*) corresponds to an atom, and each pair of directed edges represents a covalent bond in both directions.

Each atom was encoded using a **22-dimensional feature vector** comprising one-hot indicators for common atomic numbers (C, N, O, F, P, S, Cl, Br, and I), an additional “other element” indicator, scaled atomic number, scaled total degree, scaled formal charge, scaled hydrogen count, aromaticity, ring membership, one-hot hybridization state (*s p, s p*^2^, *s p*^3^, *s p*^3^*d*, and *s p*^3^*d*^2^), and a chirality-present indicator. Each bond was represented by a **6-dimensional edge vector** indicating single, double, triple, or aromatic bond type together with conjugation and ring membership. The exact atom- and bond-feature encoding, including numerical scaling, is provided in Supporting Information Table S2.

### 2.5 GPCR Sequence Representation Using Protein Language Models

GPCR sequences were indexed by UniProt accession and aligned to a deterministic receptor index before embedding generation. FASTA records and sequence metadata were cross-validated to ensure one-to-one correspondence and consistent ordering.

Three pretrained PLMs were evaluated during internal model development:

- ESM-2: `esm2_t12_35M_UR50D` [12];
- ESM-2: `esm2_t33_650M_UR50D` [12];
- ProtT5-XL-UniRef50: `prot_t5_xl_uniref50` [13]

The PLMs were used as **frozen feature extractors** rather than fine-tuned jointly with the activity-prediction network. For a GPCR sequence:

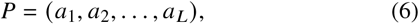

the PLM generates residue-level hidden states:

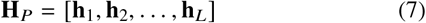

Special and padding tokens were excluded from pooling, and the receptor representation was obtained by residue-wise mean pooling:

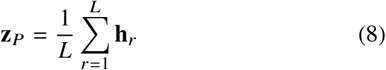

Sequences exceeding the positional capacity of a PLM were divided into nonoverlapping residue chunks. If sequence (*P*) was represented by (*K*) chunks with lengths (*L*_*k*_) and pooled chunk representations (**z**_*k*_), the final receptor vector was obtained by length-weighted aggregation:

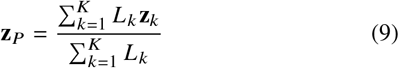

The resulting embeddings were stored as float32 arrays and reused during downstream training. Among the evaluated PLM representations, the ESM-2 650M representation was used in the selected CustomGNN model reported for the principal external and case-study analyses. Additional sequence sanitization, receptor-index quality control, and model-specific embedding-generation settings are provided in Supporting Information Section S2 and Table S3.

### 2.6 Custom Multimodal Graph Neural Network

The proposed CustomGNN consists of a ligand graph encoder, GPCR representation projection, multimodal fusion network, and two task-specific output heads.

#### 2.6.1 Initial Node and Edge Projection

Let (**x**_*v*_ ∈ ℝ^22^) denote the initial atom features of node (*v*). Atom features were first projected into a hidden representation:

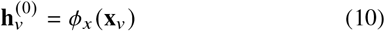

where (*ϕ*_*x*_) comprises a linear transformation followed by batch normalization, ReLU activation [33], and dropout.

Similarly, each edge vector (**e**_*uv*_ ∈ ℝ^6^) was projected into the graph hidden space:

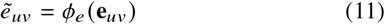

using a two-layer nonlinear edge projection.

#### 2.6.2 Residual Message Passing

For the message-passing layer (*ℓ*), an edge-conditioned message was calculated from the source-node state and projected edge features:

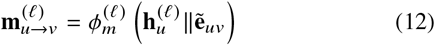

where (∥) denotes concatenation. Messages arriving at node (*v*) were summed:

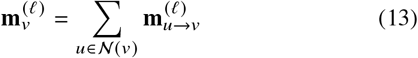

The node state was then updated using the current representation and aggregated neighborhood information:

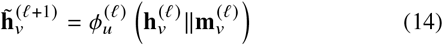

A residual connection followed by layer normalization produced:

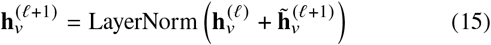

Both message and update networks comprised linear layers with batch normalization, ReLU activation, and dropout. The final selected configuration used four message-passing layers with hidden dimension 384 and dropout 0.20.

### 2.6.3 Graph-Level Readout

Following the final message-passing layer, atom embeddings were summarized using concatenated global mean and max pooling:

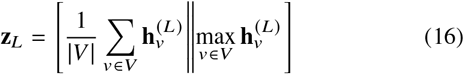

For the selected 384-dimensional graph hidden space, this produced a 768-dimensional ligand representation.

#### 2.6.4 GPCR Projection and Multimodal Fusion

The precomputed GPCR embedding was projected into the same 384-dimensional latent space using:

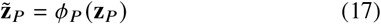

where (*ϕ*_*P*_) comprises a linear layer, layer normalization, ReLU activation, and dropout.

The ligand and receptor vectors were then concatenated:

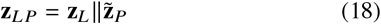

For the selected model, the 768-dimensional ligand representation and 384-dimensional receptor projection generated a 1152-dimensional multimodal vector. This vector was passed through a shared fusion network:

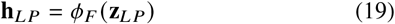

implemented as a 512-unit layer followed by a 256-unit layer, with batch normalization, ReLU activation, and dropout after each hidden transformation.

Two linear output heads were applied to the shared 256-dimensional representation:

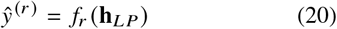

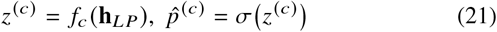

where (*ŷ*^(*r*)^) is the predicted pActivity, (*z*^(*c*)^) is the activity logit, and (*σ*) is the logistic sigmoid function [34].

The complete input representations, latent dimensions, and principal architectural components of the selected CustomGNN are summarized in Table 2.

### 2.7 Masked Multitask Objective and Optimization

Regression was optimized using the Smooth L1 (Huber) loss [35] with transition parameter (*β* = 0.5). For residual 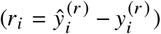,

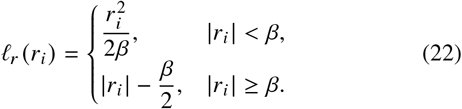

Applying the regression mask gives:

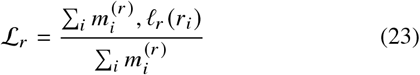

Binary classification was optimized using weighted binary cross-entropy with logits. To address the strong activity-class imbalance, the positive-class weight was estimated from the training partition as:

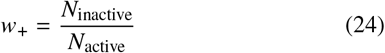

with the GNN implementation restricting extreme values to the interval ([0.05, 20]). The masked classification loss was:

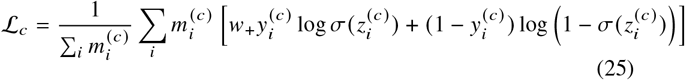

The final multitask objective was:

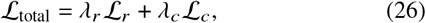

with

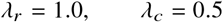

Model parameters were optimized using AdamW with a learning rate of (10^−3^), weight decay (10^−4^), and gradient-norm clipping at 5.0. A ReduceLROnPlateau scheduler reduced the learning rate by a factor of 0.5 when the validation selection criterion failed to improve.

For multitask training, checkpoints were selected by minimizing:

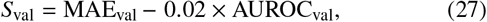

thereby retaining regression error as the dominant selection component while incorporating classification discrimination as a secondary criterion. Training was allowed to proceed for a maximum of 200 epochs with early-stopping patience of 15 epochs. The best validation checkpoint, rather than the final training epoch, was used for test evaluation.

The complete masked multitask optimization and checkpoint-selection procedure is summarized in Algorithm 2.

#### Algorithm 2

Multimodal masked multitask GNN training

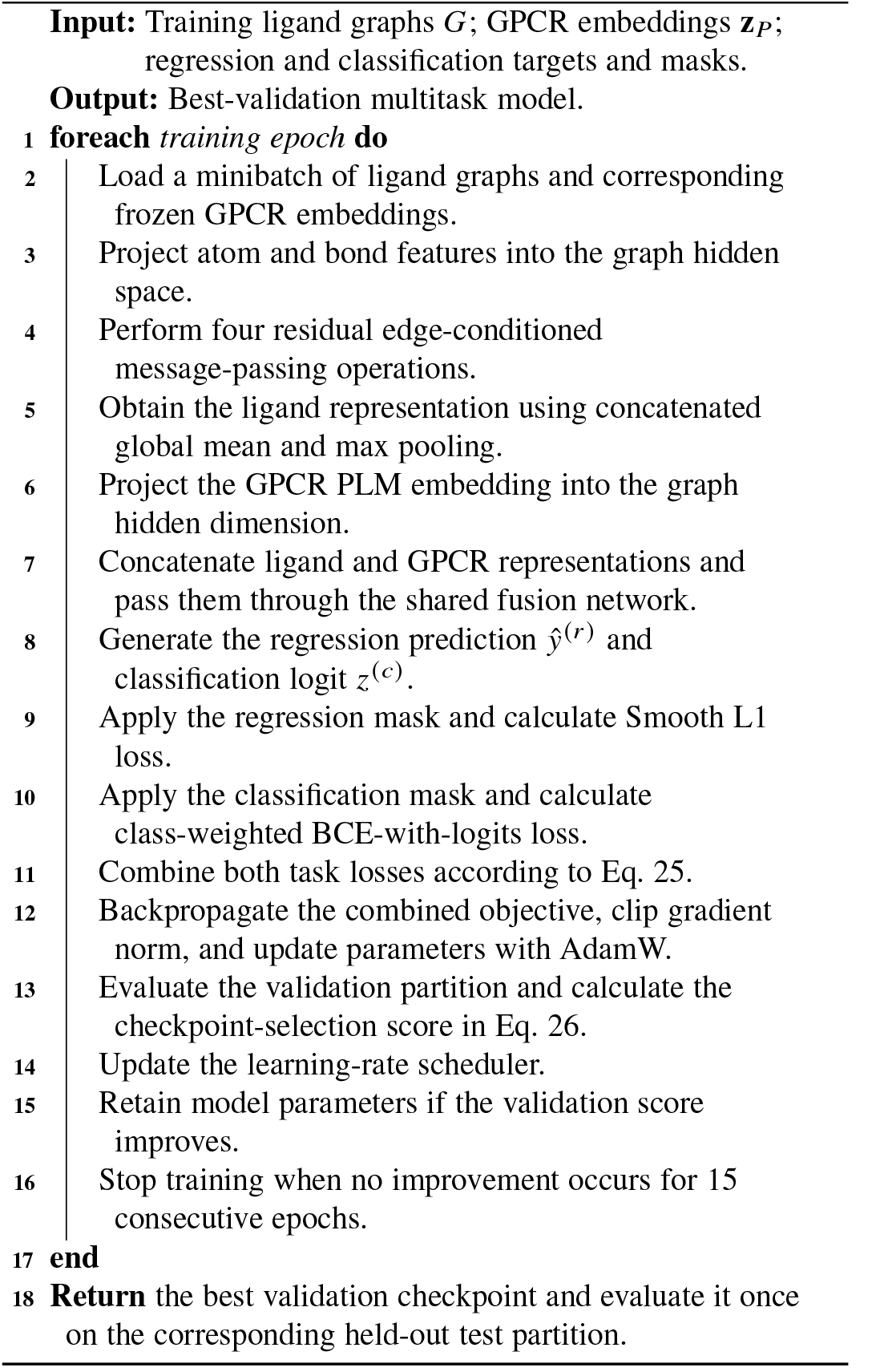

### 2.8 Dataset Partitioning and Leakage Control

The final internal dataset was partitioned into training, validation, and test subsets using a nominal 70:15:15 ratio. Three splitting schemes were used to quantify increasingly stringent forms of generalization.

#### 2.8.1 Random Pair Split

The random split was stratified using both classification state and quantitative activity. Each observation was assigned to one of three classification states, active, inactive, or ambiguous, and regression targets were divided into five quantile-based pActivity bins. The combined classification-state/pActivity-bin label was used for stratified sampling wherever sufficient samples were available. This yielded 190,217 training pairs, 40,761 validation pairs, and 40,761 test pairs.

#### 2.8.2 Bemis–Murcko Scaffold Split

Canonical ligand structures were converted to Bemis–Murcko scaffolds without chirality. All observations associated with ligands sharing the same scaffold were assigned to the same partition. Scaffold groups were ordered by decreasing group size and allocated greedily to the partition with the smallest occupancy relative to its target size. This prevented a molecular scaffold from occurring in more than one split while maintaining approximately 70:15:15 pair counts.

The scaffold split contained 190,217 training pairs, 40,761 validation pairs, and 40,761 test pairs, corresponding to 128,353, 27,664, and 27,677 unique ligands, respectively.

### 2.8.3 Cold-Ligand Split

For the strict cold-ligand evaluation, the standard InChIKey served as the grouping variable. Every occurrence of a ligand, irrespective of GPCR target, was therefore confined to exactly one of the training, validation, or test partitions. Ligand groups were allocated using the same group-size-aware greedy procedure. The resulting split contained 128,588 unique training ligands and 27,553 unique ligands in each of the validation and test partitions, with the same pair counts as the other splitting strategies.

All three primary splits retained all 216 GPCR targets. Automated leakage checks verified group uniqueness across partitions for the grouped splitting schemes. Complete partition composition and leakage-control criteria are reported in Supporting Information Table S4.

### 2.9 Hyperparameter Optimization and Final Model Training

Hyperparameter optimization was conducted on the internal data before final multi-seed evaluation. Seven candidate configurations were assessed by varying graph hidden dimension, number of message-passing layers, dropout, and learning rate:

- baseline: 256 hidden units, 4 layers, dropout 0.20, learning rate 10^−3^;
- lower learning rate: 256 units, 4 layers, dropout 0.20, learning rate 5 × 10^−4^;
- low dropout: 256 units, 4 layers, dropout 0.10;
- high dropout: 256 units, 4 layers, dropout 0.30;
- shallow: 256 units, 3 message-passing layers;
- deeper: 256 units, 5 message-passing layers;
- wider: 384 units, 4 message-passing layers, dropout 0.20.

HPO experiments were performed using the random, scaffold, and cold-ligand splits and the evaluated protein representations. Candidate configurations were ranked according to mean validation classification AUROC from the HPO result summaries. The wider configuration (384 hidden units, four graph layers, dropout 0.20, learning rate (10^−3^) was selected.

Final GNN experiments were repeated with three independent random seeds:

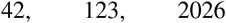

The final training protocol used a batch size of 512 for optimization and 768 for validation/test inference, a maximum of 200 epochs, early-stopping patience of 15, weight decay (10^−4^), regression and classification loss weights of 1.0 and 0.5, respectively, and the HPO-selected architecture. ESM-2 35M, ESM-2 650M, and ProtT5 receptor representations were evaluated under this common protocol. The complete HPO experimental design and selection settings are summarized in Supporting Information Table S5.

### 2.10 MLP Baseline

A multitask neural MLP was used as the principal baseline to determine whether explicitly learned molecular-graph representations provided an advantage over fixed cheminformatics features.

The protein-aware MLP input was constructed as:

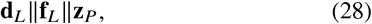

where (**d**_*L*_) denotes the RDKit descriptor vector, (**f**_*L*_) the 2048-bit radius-2 Morgan fingerprint, and (**z**_*P*_) the corresponding GPCR PLM representation. The comparison used the ESM-2 650M receptor representation for consistency with the selected graph model.

The shared MLP trunk used hidden dimensions:

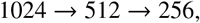

with batch normalization, GELU activation [36], and dropout of 0.25. Separate task-specific regression and classification heads branched from the final shared representation. Both heads contained an additional hidden transformation followed by the corresponding scalar output.

Feature normalization and regression-target standardization were fitted exclusively on each training partition. The MLP used the same conceptual masked multitask formulation as the GNN, with Smooth L1 regression loss (*β* = 0.5) and class-weighted BCE-with-logits classification loss. Optimization used AdamW, learning rate (10^−3^), weight decay (10^−4^), and batch size 2048. The MLP was evaluated only in the internal model-comparison experiments and was not evaluated on the external test set. Additional architectural, optimization, and validation-threshold details for the MLP baseline are provided in Supporting Information Table S7.

### 2.11 Architectural Reference and Single-Task Ablations

A second graph implementation based on the Graph Isomorphism Network with Edge features (GINE) in PyTorch Geometric was trained using matched graph inputs, protein embeddings, split definitions, optimization settings, and HPO-selected hyperparameters. This implementation served as an architectural reference for assessing whether results were specific to the custom message-passing formulation; it was not treated as the principal baseline. The complete PyTorch Geometric GINE reference architecture is provided in Supporting Information Table S6.

To isolate the contribution of joint multitask optimization, the selected CustomGNN with ESM-2 650M receptor embeddings was additionally trained in two single-task modes:

- **regression-only**, in which only the Smooth L1 regression loss contributed gradients; and
- **classification-only**, in which only the weighted binary classification loss contributed gradients.

The network architecture and both output heads were retained in all modes, but the irrelevant task loss was omitted from gradient calculation. Regression-only checkpoints were selected using validation MAE, whereas classification-only checkpoints were selected using validation Matthews correlation coefficient (MCC) [37]. Ablations were performed for all three primary split strategies and all three random seeds.

### 2.12 Performance Evaluation

Regression performance was evaluated using mean absolute error (MAE), root-mean-square error (RMSE), and coefficient of determination (*R*^2^). Pearson (*R*) and Spearman (*ρ*) correlations were additionally calculated for analyses in which correlation structure was of interest, including the MLP and selectivity analyses.

For (*N*) valid regression observations:

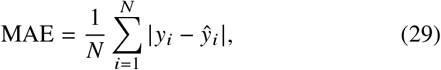

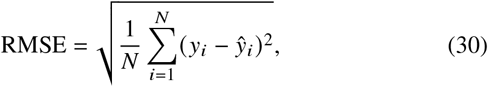

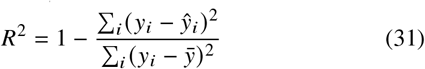

Classification evaluation included accuracy, balanced accuracy, precision, recall, and F1 score for both active and inactive classes, MCC, receiver-operating-characteristic area under the curve (ROC-AUC), and precision–recall area under the curve (PR-AUC) for both the active and inactive classes. Given the marked activity-class imbalance, balanced and class-specific metrics were retained alongside conventional accuracy and ROC-AUC.

For the CustomGNN, class predictions used a probability threshold of 0.5 after sigmoid transformation of the classification logit. The MLP baseline selected its classification threshold from the validation partition by maximizing MCC over candidate thresholds from 0.05 to 0.95, and this fixed validation-derived threshold was subsequently applied to the corresponding test partition. Test-set observations were never used for threshold selection, hyperparameter optimization, or early stopping.

Final internally reported GNN performance was summarized across the three independent seeds, while individual-seed outputs were retained for reproducibility and supplementary analyses. The complete experimental design, final optimization settings, and evaluation protocol are consolidated in Table 3.

### 2.13 Independent External Evaluation

An independent external evaluation set was constructed using ChEMBL release 37 together with an independently filtered BindingDB [6] component. The same GPCR, activity-type, assay-confidence, pActivity, molecular-standardization, and multitask-label definitions used for the internal dataset were retained wherever applicable.

For the ChEMBL component, activity identifiers present in ChEMBL 37 but absent from the ChEMBL 36 extraction were first identified, thereby defining the release-delta activity pool. These records were subsequently processed through the same activity-standardization and ligand–target aggregation workflow used for the training data.

For BindingDB, records were restricted to human GPCR targets represented by a unique mapped UniProt accession. Rows lacking molecular structures were discarded; modified or explicitly mutant/chimeric targets were excluded by default, and publication-associated records were required. BindingDB rows explicitly originating from ChEMBL were removed to avoid duplicating the ChEMBL release-delta component. Eligible (*K*_*i*_), (*K*_*d*_), (*IC*_50_), and (*EC*_50_) measurements were converted to the common pActivity framework.

To prevent training-pair leakage, molecular structures from the frozen training set were re-standardized and used to construct reference keys combining GPCR UniProt accession with standardized parent-ligand InChIKey. External candidates were annotated for pair, ligand, connectivity, scaffold, and target novelty relative to the ChEMBL 36 training data. Any ligand–target pair already represented in the internal training reference was excluded from the strict external set. BindingDB pairs already represented in the ChEMBL 37 release-delta component were also removed before source combination.

A second leakage audit was performed after external feature generation. Detection of any standardized ligand–target key present in the frozen training reference caused evaluation-set finalization to terminate rather than silently retain the overlapping pair. The external-set audit criteria, final composition, and structural/target novelty definitions are summarized in Supporting Information Section S5 and Table S8.

The finalized external test set contained 6,319 ligand–GPCR pairs, 3,421 unique ligands, and 159 GPCR targets. All 6,319 observations were valid for regression, while 5,021 had valid binary labels, comprising 4,834 active and 187 inactive interactions. No class balancing or external-data resampling was applied.

The external set was used only after completion of internal model development. GNN configurations were evaluated without retraining on the external observations. The principal external results presented in the manuscript correspond to the selected CustomGNN model, whereas the MLP baseline was restricted to internal evaluation.

### 2.14 Held-Out DRD2–DRD3 Selectivity Case Study

A receptor-subtype selectivity analysis was constructed for DRD2 and DRD3. Ligands were required to have compatible quantitative measurements for both receptors. Mixed endpoint-type combinations were excluded by default so that each matched comparison was based on a common activity-type group.

Where multiple eligible measurements remained for a ligand–receptor/activity-type combination, receptor-specific activity was aggregated by the median. Rows with an aggregate pActivity standard deviation greater than 1.0 were excluded from the case-study construction.

For each matched ligand, observed selectivity was defined as:

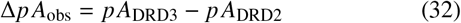

Accordingly, (Δ*p A*_obs_ > 0) indicates greater measured activity toward DRD3, whereas (Δ*p A*_obs_ < 0) indicates greater measured activity toward DRD2. A value of (Δ*p A*_obs_ = 0) denotes equivalent measured activity toward the two receptors. Strong receptor selectivity was defined as:

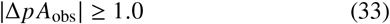

corresponding to an approximately tenfold or greater difference in activity between DRD2 and DRD3 on the logarithmic pActivity scale.

The assembled case-study resource contained 4,527 matched ligands with measurements for both DRD2 and DRD3, of which 2,410 satisfied (|Δ*p A*| ≥ 1.0). The dataset included 4,431 internally derived ChEMBL 36 pairs and 96 externally derived ChEMBL 37/BindingDB pairs. Before case-study inference, the internal cohort was cross-referenced against the predefined split manifests, and only compounds satisfying the held-out test-partition criteria were retained for analysis. Additional cohort-composition and held-out quality-control information is provided in Supporting Information Section S6 and Table S9.

For model predictions, receptor-specific predicted activities were combined analogously:

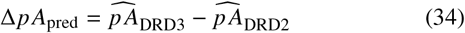

Predictions from the three independently trained seeds were treated as repeated model realizations rather than independent compounds. The primary selectivity estimate for each ligand was therefore the mean predicted (Δ*p A*_pred_) across seeds. Performance was evaluated separately for each splitting strategy to avoid double-counting ligands that could occur in the test subsets of more than one split definition.

Selectivity regression was assessed using RMSE, MAE, (*R*^2^), Spearman correlation, and Pearson correlation. Bootstrap 95% confidence intervals were calculated over unique ligand pairs using 1,000 bootstrap resamples. Seed-wise metric mean and standard deviation were retained as secondary measures of model variability. Classification-oriented analyses additionally evaluated recovery of receptor-preference/selectivity categories derived from the observed (Δ*p A*).

### 2.15 Software and Computational Environment

The computational pipeline was implemented in Python 3.11.3 and executed in a Linux-based high-performance computing environment using NVIDIA A100-series GPUs with CUDA 12.0.0. Neural-network modelling used PyTorch 2.1.2 [38] and PyTorch Geometric 2.5.0. Protein language-model inference used the Hugging Face `transformers` [39] framework (v4.30+). Molecular processing, standardization, descriptor generation, fingerprinting, scaffold calculation, and graph construction were performed using RDKit 2024.03.3. Conventional preprocessing and metric calculations used scikit-learn 1.3.1 [40], and manuscript visualizations were generated using Matplotlib 3.7.2 [41].

Exact molecular feature definitions, PLM preprocessing and index-validation procedures, internal split composition and leakage controls, reference-model and baseline configurations, external-set audit details, and DRD2-DRD3 cohort quality-control information are provided in the Supporting Information.

## 3 Results and Discussion

### 3.1 Dataset Characteristics

The curated ChEMBL 36 dataset comprised 271,739 lig- and–GPCR pairs, spanning 183,694 unique ligands and 216 human GPCR targets. All observations carried a quantitative *p Activit y* value and were retained for regression. Binary activity labels were assigned to 225,527 pairs after masking the intermediate 5 < *p Activit y* < 6 interval; 212,492 of these pairs were active and 13,035 inactive, producing a markedly imbalanced classification problem (94.2% versus 5.8%). The remaining 46,212 intermediate-activity observations contributed to regression but not to the classification loss (Table 1).

**Table 1:** Dataset construction and multitask target definition.

| Component | Criterion or definition | Final setting/value |
| --- | --- | --- |
| Internal bioactivity source | ChEMBL | Release 36 |
| Organism | Target organism | *Homo sapiens* |
| Target type | ChEMBL target type | Single protein |
| Target family | Protein-class/target annotation | GPCR |
| Quantitative endpoints | Accepted activity types | $K_i$, $K_d$, $IC_{50}$, $EC_{50}$ |
| Assay confidence | ChEMBL confidence score | $\geq 7$ |
| Activity scale | Standardized target | $pA = -\log_{10}(A_M)$ |
| Accepted pActivity range | QC interval | 2–12 |
| Replicate aggregation | Ligand–GPCR level | Median pActivity |
| Active label | Binary classification | $pA \geq 6.0$ |
| Inactive label | Binary classification | $pA \leq 5.0$ |
| Ambiguous interval | Classification masked | $5.0 < pA < 6.0$ |
| Minimum GPCR support | Pairs per receptor | 50 |
| Final ligand–GPCR pairs | — | 271,739 |
| Unique ligands | — | 183,694 |
| GPCR targets | — | 216 |
| Regression-valid observations | — | 271,739 |
| Classification-valid observations | — | 225,527 |
| Active / inactive | — | 212,492 / 13,035 |

**Table 2:**
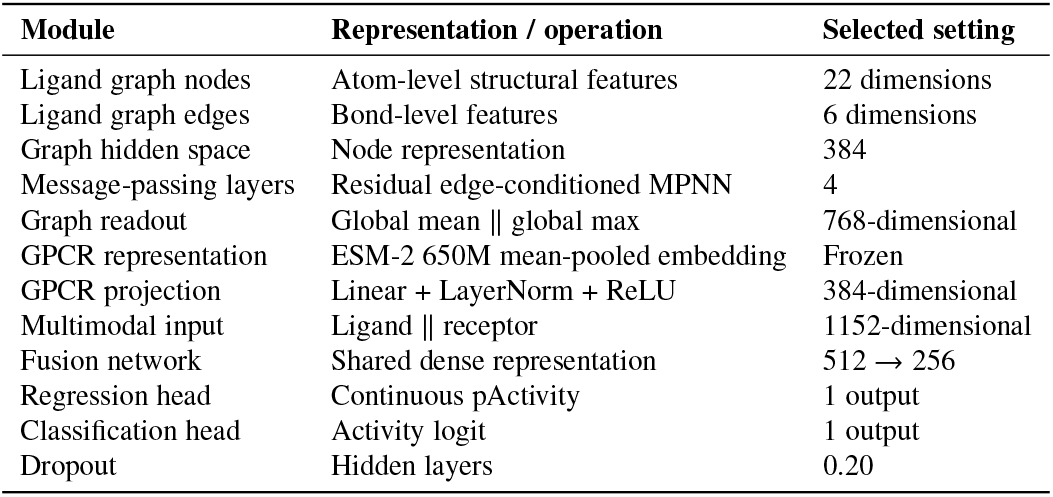
Input representations and selected CustomGNN architecture.

**Table 3:** Experimental design and optimization protocol.

| Category | Parameter | Setting |
| --- | --- | --- |
| Internal splits | Strategies | Random, scaffold, cold-ligand |
| Partition ratio | Train/validation/test | 70/15/15 |
| Split seed | Initial split generation | 42 |
| Final model seeds | Repeated training | 42, 123, 2026 |
| Selected graph hidden dimension | HPO result | 384 |
| Message-passing layers | HPO result | 4 |
| Dropout | HPO result | 0.20 |
| Fusion dimensions | Shared network | 512 → 256 |
| Optimizer | — | AdamW |
| Learning rate | — | $1 \times 10^{-3}$ |
| Weight decay | — | $1 \times 10^{-4}$ |
| GNN train batch | — | 512 |
| GNN evaluation batch | — | 768 |
| Maximum epochs | — | 200 |
| Early-stopping patience | — | 15 |
| Regression loss | — | Smooth L1, $\beta = 0.5$ |
| Classification loss | — | weighted BCE-with-logits |
| Regression/classification weights | — | 1.0 / 0.5 |
| Gradient clipping | Maximum norm | 5.0 |
| Primary regression metrics | — | MAE, RMSE, $R^2$ |
| Classification metrics | — | ROC-AUC, PR-AUC, MCC, balanced accuracy, precision, recall, F1 |

Random, Bemis–Murcko scaffold, and strict cold-ligand partitioning each produced 190,217 training, 40,761 validation, and 40,761 test observations while retaining all 216 receptors. The comparison among these splits therefore isolates increasingly stringent forms of chemical generalization without introducing differences in target coverage. The scaffold split excluded shared Bemis–Murcko frameworks across partitions, whereas the cold-ligand split excluded exact ligand identity irrespective of target. This distinction becomes consequential below: excluding complete scaffolds produced a substantially larger loss of predictive accuracy than excluding exact ligands alone.

The independent external set comprised 6,319 pairs from 3,421 ligands and 159 GPCRs, of which 5,021 were classification-valid. Its class composition was even more asymmetric than that of the internal data, with 4,834 active and only 187 inactive observations. Consequently, accuracy and positive-class precision–recall metrics were interpreted alongside MCC, balanced accuracy, and inactive-class performance throughout. The DRD2–DRD3 selectivity resource comprised a separate matched cohort of 4,527 compounds measured against both receptors, including 2,410 compounds with |Δ*p Activit y*| ≥ 1.

### 3.2 Model Selection

Hyperparameter optimization identified a broader latent graph space, rather than greater depth or stronger regularization, as the most favourable modification to the base architecture. Across the nine combinations of internal split and protein representation, the 384-dimensional, four-layer model achieved a mean validation ROC-AUC of 0.9591, compared with 0.9587 for the 256-dimensional baseline and 0.9582 for the lower-learning-rate configuration. Deeper, shallower, and altered-dropout variants were slightly weaker. The absolute separation among the leading configurations was therefore small, arguing against a fragile hyper-parameter optimum. The 384-dimensional architecture was fixed before the final three-seed evaluation.

The three pretrained receptor representations: ESM-2 35M, ESM-2 650M, and ProtT5, subsequently occupied a narrow performance range. For example, random-split CustomGNN MAE varied only from 0.513 to 0.514 across the three PLMs, and scaffold MAE from 0.641 to 0.646. No representation dominated every split or endpoint: ProtT5 was marginally strongest on selected classification measures, ESM-2 35M slightly favoured individual cold-ligand results, and ESM-2 650M gave the lowest mean scaffold MAE while remaining effectively tied with the best model under random evaluation. ESM-2 650M was therefore retained as the principal receptor representation for downstream analyses, while the full PLM comparison was preserved rather than interpreting these marginal differences as evidence of model-scale superiority.

### 3.3 Internal Generalization

Chemical scaffold novelty, rather than ligand identity alone, defined the principal internal generalization barrier. The selected CustomGNN achieved an MAE of 0.513 ± 0.006 under random splitting and retained most of this accuracy when evaluated on completely unseen ligands (0.540 ± 0.006). In contrast, withholding Bemis–Murcko scaffolds increased MAE to 0.641 ± 0.005 and reduced *R*^2^ from 0.695 ± 0.005 to 0.546 ± 0.005 (Table 4; Fig. 1A, 1B, 1E). Exact ligand exclusion therefore incurred only a 0.027-unit MAE penalty relative to random evaluation, whereas chemotype-level exclusion increased error by 0.128 units.

**Table 4:** Internal performance of the principal ESM-2 650M models. GNN values are reported as mean ± SD over seeds 42, 123, and 2026. The MLP baseline was evaluated once for each split.

| Model | Split | MAE | RMSE | R<sup>2</sup> | ROC-AUC | MCC | Balanced accuracy |
| --- | --- | --- | --- | --- | --- | --- | --- |
| CustomGNN | Random | **0.513 ± 0.006** | **0.704 ± 0.006** | **0.695 ± 0.005** | **0.971 ± 0.000** | **0.661 ± 0.020** | **0.913 ± 0.003** |
| PyG-GINE | Random | 0.565 ± 0.006 | 0.760 ± 0.007 | 0.645 ± 0.007 | 0.969 ± 0.002 | 0.532 ± 0.017 | 0.910 ± 0.006 |
| MLP | Random | 0.851 | 1.079 | 0.283 | 0.735 | 0.240 | 0.730 |
| CustomGNN | Scaffold | **0.641 ± 0.005** | **0.858 ± 0.004** | **0.546 ± 0.005** | **0.941 ± 0.001** | **0.529 ± 0.022** | **0.862 ± 0.004** |
| PyG-GINE | Scaffold | 0.690 ± 0.014 | 0.916 ± 0.013 | 0.481 ± 0.015 | 0.936 ± 0.008 | 0.435 ± 0.028 | 0.861 ± 0.010 |
| MLP | Scaffold | 0.874 | 1.103 | 0.249 | 0.730 | 0.241 | 0.726 |
| CustomGNN | Cold ligand | **0.540 ± 0.006** | **0.736 ± 0.006** | **0.670 ± 0.005** | **0.965 ± 0.003** | **0.626 ± 0.016** | 0.893 ± 0.003 |
| PyG-GINE | Cold ligand | 0.593 ± 0.010 | 0.790 ± 0.011 | 0.619 ± 0.011 | 0.961 ± 0.001 | 0.520 ± 0.017 | **0.899 ± 0.004** |
| MLP | Cold ligand | 0.859 | 1.085 | 0.282 | 0.760 | 0.279 | 0.755 |

**Fig. 1:**
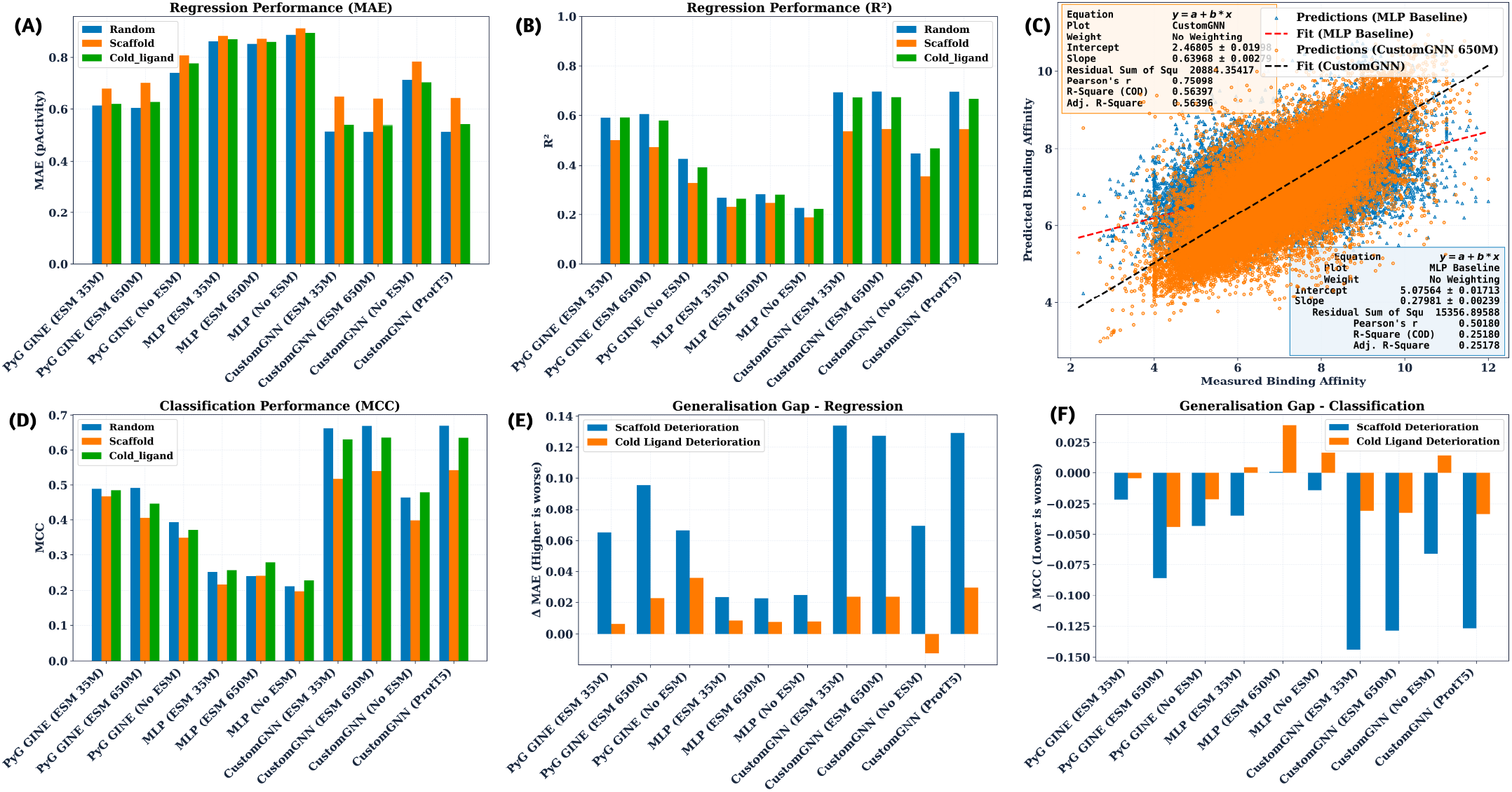
Internal predictive performance and generalization across chemical split strategies. (A) Test MAE and (B) *R*^2^ for the MLP baseline, PyG-GINE, and CustomGNN across random, scaffold, and cold-ligand splits using the indicated GPCR representations. GNN values summarize seeds 42, 123, and 2026. (C) Measured versus predicted pActivity on the scaffold test set for the ESM-2 650M MLP and CustomGNN (seed 42); dashed lines indicate linear fits. (D) Matthews correlation coefficient (MCC) for activity classification. (E) Regression generalization gap, defined as ΔMAE = MAE_scaffold/cold_ − MAE_random_. (F) Classification generalization gap, defined as ΔMCC = MCC_scaffold/cold_ − MCC_random_. Error bars represent across-seed variability where shown.

This hierarchy indicates that the model transfers efficiently to unseen molecules when related structural series remain represented, but extrapolation across distinct chemical frameworks remains more difficult. That distinction is consistent with systematic molecular-property benchmarking showing that random splits can conceal limitations in chemical-space transfer and that scaffold structure and activity cliffs are major determinants of apparent model generalizability [11].

The learned molecular graph representation also produced a large advantage over the protein-aware MLP. Relative to the ESM-2 650M MLP, CustomGNN reduced MAE by 39.8% under random, 26.7% under scaffold, and 37.1% under cold-ligand evaluation. The increase in explained variance was similarly substantial: MLP *R*^2^ remained between 0.249 and 0.283, whereas CustomGNN reached 0.546–0.695. The measured-versus-predicted comparison in Fig. 1C exposes the qualitative origin of this difference: MLP predictions are strongly contracted toward the centre of the activity range, whereas CustomGNN retains substantially more of the measured variation. Split-resolved comparisons in Fig. S4 reproduce this pattern across all three test regimes.

This result is not a foregone consequence of replacing fingerprints with a graph network. Large-scale benchmarking has shown that fixed cheminformatics representations remain competitive for many molecular-property tasks and that representation learning benefits depend strongly on dataset size and chemical-space structure [11]. Here, combining a large ligand––target dataset with receptor-conditioned prediction appears to provide enough information for learned graph representations to decisively exceed the descriptor/fingerprint baseline.

Classification showed the same general ordering, but the severe class imbalance altered which metrics were informative. CustomGNN maintained ROC-AUC values of 0.971, 0.965, and 0.941 under random, cold-ligand, and scaffold evaluation, respectively, while MCC fell from 0.661 to 0.626 and 0.529 as chemical novelty increased (Fig. 1D, 1F). The corresponding inactive-class PR-AUC values were 0.811, 0.779, and 0.641, whereas active-class PR-AUC exceeded 0.995 throughout (Fig. S1). Given that 94.2% of classification-valid observations were active, the latter is partly prevalence-driven; MCC, balanced accuracy, and inactive-class recovery provide the more stringent evidence that classification performance persists beyond the majority class.

Against the MLP baseline, the advantage was unambiguous across all three independent CustomGNN seeds. Paired error comparisons showed lower MAE and MSE in every split-by-seed experiment, while bootstrap comparisons showed markedly higher ROC-AUC and active-class PR-AUC (all *p* < 0.001; Table S10). Thus, the difference between the graph model and fixed-feature baseline was substantially larger than the seed-to-seed variation of the graph model itself.

### 3.4 Receptor Representation and Graph Architecture

The largest representation-level effect was the presence of receptor information, not the identity of the protein language model. Removing the GPCR embedding increased CustomGNN MAE from 0.513 to 0.714 under random evaluation, from 0.540 to 0.702 under cold-ligand evaluation, and from 0.641 to 0.784 under scaffold evaluation (Figs. 1A, 1B and 2A, 2C). MCC deteriorated in parallel, from 0.661 to 0.465, from 0.626 to 0.479, and from 0.529 to 0.399, respectively. The receptor-independent model therefore retained substantial ligand-side signal but could not reproduce the full ligand–target relationship.

By contrast, ESM-2 35M, ESM-2 650M, and ProtT5 were nearly interchangeable at the scale of the principal performance differences (Fig. 2A–2D). Mean random-split MAEs were 0.514, 0.513, and 0.513, respectively; scaffold values were 0.646, 0.641, and 0.643. This compression across independently pretrained sequence models suggests that each encodes sufficient global receptor information for the present task and that the limiting representation problem lies elsewhere; most plausibly in how global receptor information is coupled to ligand structure rather than in simply increasing PLM size.

**Fig. 2:**
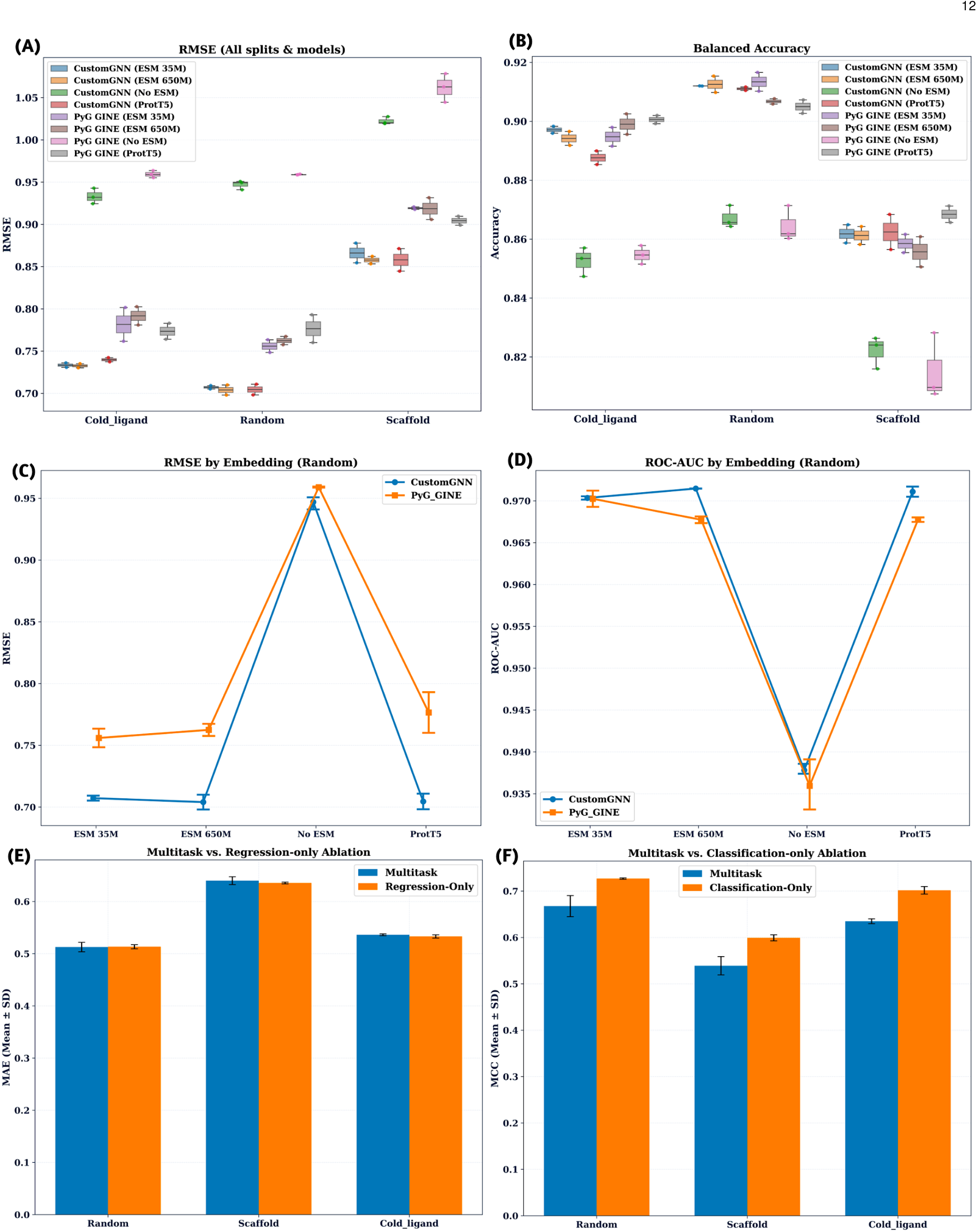
Internal predictive performance and generalization across chemical split strategies. (A) Test MAE and (B) *R*^2^ for the MLP baseline, PyG-GINE, and CustomGNN across random, scaffold, and cold-ligand splits using the indicated GPCR representations. GNN values summarize seeds 42, 123, and 2026. (C) Measured versus predicted pActivity on the scaffold test set for the ESM-2 650M MLP and CustomGNN (seed 42); dashed lines indicate linear fits. (D) Matthews correlation coefficient (MCC) for activity classification. (E) Regression generalization gap, defined as ΔMAE = MAE_scaffold/cold_ − MAE_random_. (F) Classification generalization gap, defined as ΔMCC = MCC_scaffold/cold_ − MCC_random_. Error bars represent across-seed variability where shown.

Recent structure-free protein–ligand models point in the same direction. G-PLIP combines biological and chemical representations without requiring an experimental protein–ligand complex, while PSICHIC demonstrated that sequence-derived protein information can support both affinity prediction and ligand–protein selectivity analysis. PMMR similarly used pretrained target and drug representations to improve cold-start transfer [15], [16], [17]. The present ablation provides a particularly direct test of this principle because ligand architecture, dataset partitions, and training objective were held constant while the receptor representation was removed.

The CustomGNN–GINE comparison further separated the contribution of target encoding from that of molecular message passing. Under matched ligand features, protein embeddings, data partitions, and training protocol, CustomGNN reduced ESM-2 650M MAE from 0.565 to 0.513 under random splitting, 0.593 to 0.540 under cold-ligand splitting, and 0.690 to 0.641 under scaffold splitting (Table 4; Fig. 2A, 2C). The corresponding *R*^2^ gains were 0.050, 0.051, and 0.064.

These regression gains were unusually consistent: all nine seed-wise CustomGNN–GINE MAE and MSE comparisons were significant, with paired Wilcoxon *p*-values ranging from 9.1 × 10^−66^ to below numerical precision (Table S10). The paired RMSE analysis in Fig. S2 shows the same directional shift across seeds. In contrast, ROC-AUC and PR-AUC differences were much smaller and varied by seed. The architecture therefore appears to improve the precision of quantitative pActivity estimation more strongly than global active–inactive ranking.

MCC nevertheless increased materially, from 0.532 to 0.661 under random, 0.520 to 0.626 under cold-ligand, and 0.435 to 0.529 under scaffold evaluation, indicating improved threshold-level separation despite relatively small changes in ROC-AUC. The custom edge-conditioned residual message-passing formulation therefore conferred a reproducible regression advantage and a more moderate classification advantage over the GINE reference, rather than a uniform shift across every endpoint.

Training trajectories reinforce this interpretation. Validation MAE and MCC followed similar convergence patterns across seeds and split strategies (Fig. S3), while the final regression standard deviations were typically < 0.01*p Activit y* units. The principal architectural conclusions are therefore robust to initialization rather than being driven by a single favourable training trajectory.

### 3.5 Multitask Effects

Joint learning did not simply improve the optimum of each task; it produced a balanced shared representation with a measurable trade-off between regression and classification objectives (Fig. 2E, 2F).

Regression-only training was essentially indistinguishable from multitask training under random splitting (MAE 0.513 versus 0.513) and provided only small gains under scaffold and cold-ligand evaluation, where MAE decreased from 0.641 to 0.636 and from 0.540 to 0.533, respectively. The classification objective therefore imposed little cost on quantitative *p Activit y* prediction.

The converse comparison was more metric-dependent. Classification-only training increased MCC from 0.661 to 0.727 under random evaluation, from 0.529 to 0.599 under scaffold evaluation, and from 0.626 to 0.702 under cold-ligand evaluation. Yet ROC-AUC decreased from 0.971 to 0.960, from 0.941 to 0.931, and from 0.965 to 0.943; balanced accuracy also declined in all three settings. Classification-only optimization therefore sharpened the particular decision boundary reflected by MCC while weakening global probability ranking.

The relevant conclusion is not that multitask learning universally dominates single-task optimization. Rather, it provides a single model that preserves regression performance near the regression-specific optimum while retaining strong classification ranking and threshold discrimination. This is the more useful property for a ligand-prioritization model in which continuous potency estimation and active/inactive assessment are required simultaneously.

Recent work has independently explored this coupling. SLAB jointly optimizes affinity regression and activity classification rather than learning the two endpoints in isolation and reported gains from sharing representations between the tasks [20]. The present results add an important qualification: positive transfer is metric-dependent, and a fixed multitask weighting need not sit at the optimum for either task individually. Adaptive task weighting or gradient-balancing strategies therefore represent a plausible route to improving the present architecture without changing its underlying ligand–receptor representation.

### 3.6 External Generalization

Independent bioactivity data exposed a domain shift that none of the internal split strategies fully anticipated. On the 6,319-pair ChEMBL 37/BindingDB test set, the random-trained ESM-2 650M CustomGNN reached an MAE of 0.949 ± 0.007 and *R*^2^ = 0.093 ± 0.010, compared with 0.513 and 0.695 internally. Cold-ligand and scaffold training did not materially close this gap: external MAE remained 0.943 ± 0.012 and 0.953 ± 0.005, respectively, with *R*^2^ near 0.10 in both cases (Table 5).

**Table 5:** Performance on the independent ChEMBL 37/BindingDB evaluation set. Values are mean ± SD over the three independently trained ESM-2 650M CustomGNN models.

| Training split | MAE | RMSE | R<sup>2</sup> | ROC-AUC | MCC | Balanced accuracy | Inactive PR-AUC |
| --- | --- | --- | --- | --- | --- | --- | --- |
| Random | 0.949 ± 0.007 | 1.245 ± 0.007 | 0.093 ± 0.010 | 0.765 ± 0.015 | **0.171 ± 0.022** | **0.666 ± 0.036** | 0.128 ± 0.024 |
| Cold ligand | **0.943 ± 0.012** | **1.237 ± 0.020** | **0.104 ± 0.029** | 0.737 ± 0.021 | 0.140 ± 0.027 | 0.635 ± 0.029 | **0.129 ± 0.023** |
| Scaffold | 0.953 ± 0.005 | 1.240 ± 0.008 | 0.101 ± 0.012 | **0.765 ± 0.006** | 0.153 ± 0.026 | 0.659 ± 0.034 | 0.109 ± 0.010 |

The external measured-versus-predicted distributions reveal a systematic contraction of the prediction range (Fig. 3A–3C). Pearson correlations remained positive (*r* ≈ 0.46–0.48), but high-pActivity observations were underpredicted and low-pActivity observations shifted upward. Thus, the model retained useful ordering while losing calibration and variance under domain shift. This distinction explains the coexistence of moderate correlations with *R*^2^ values close to 0.1.

**Fig. 3:**
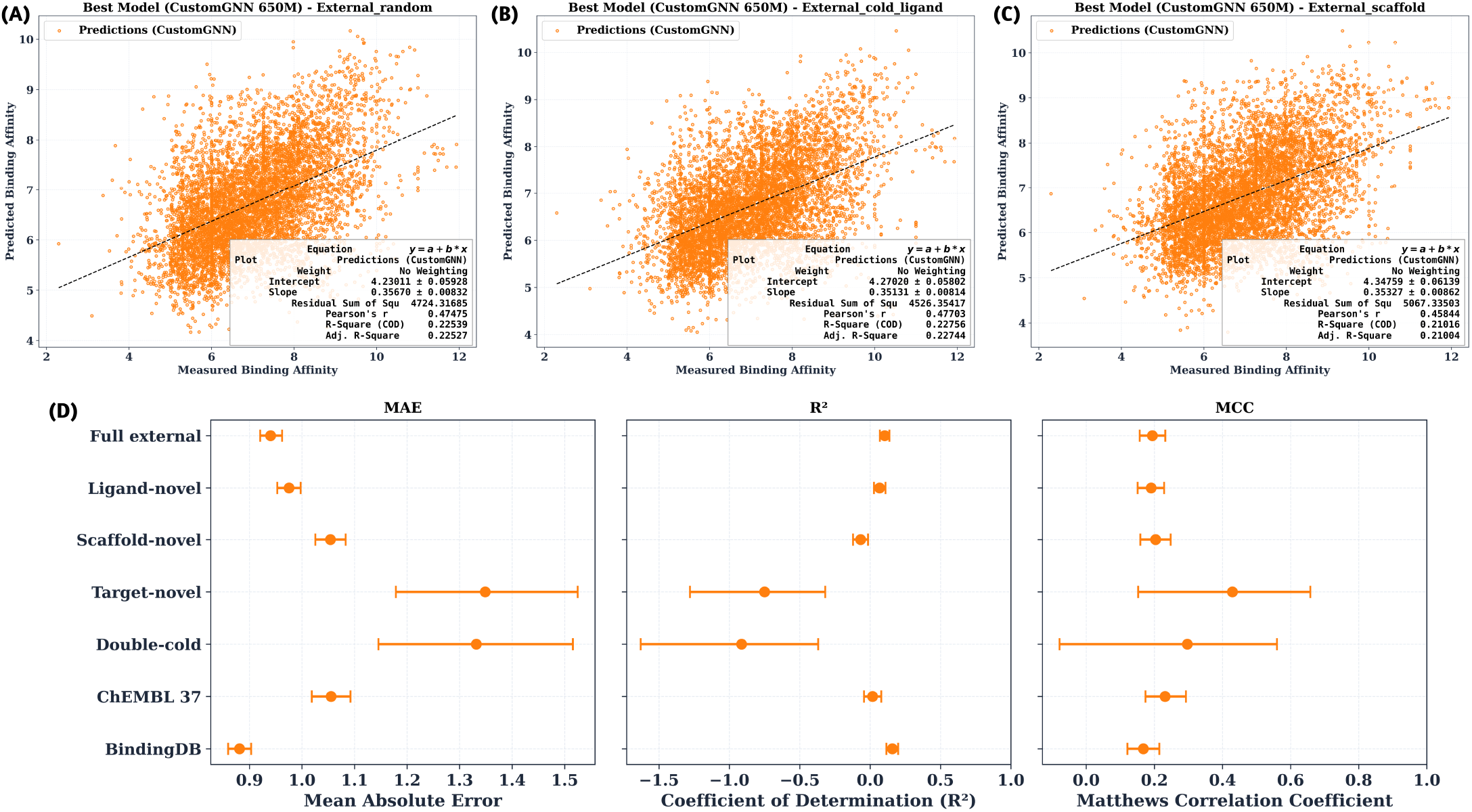
Independent external evaluation of the selected CustomGNN. (A–C) Measured versus predicted pActivity on the independent ChEMBL 37/BindingDB evaluation set for ESM-2 650M CustomGNN models trained using the (A) random, (B) cold-ligand, and (C) scaffold splits. Dashed lines indicate linear fits. (D) MAE, *R*^2^, and MCC across the full external set and ligand-novel, scaffold-novel, target-novel, double-cold, ChEMBL 37, and BindingDB subsets. Horizontal error bars denote 95% bootstrap confidence intervals. Models were evaluated without retraining on the external data.

The near-convergence of the random, cold-ligand, and scaffold-trained models on the same external performance is itself informative. Scaffold-disjoint evaluation is considerably harder than random evaluation internally, yet both internal regimes substantially underestimate the difficulty of later-release and independently sourced activity data. Chemical disjointness is therefore necessary for credible internal validation, but it is not equivalent to temporal/source generalization.

This observation parallels recent DTI work in which performance degrades as evaluation moves from transductive to unseen-drug and unseen-target regimes. ZeroBind explicitly framed novel drug/protein prediction as a zero-shot generalization problem, while PMMR used pretrained multimodal representations to address loss of accuracy outside the training domain [17], [22]. The present external test adds another axis: even when targets or chemical motifs overlap partially with the training domain, release-to-release and database-to-database variation introduces a substantial additional shift.

Novelty stratification resolved part of that shift (Fig. 3D). Ligand-novel observations incurred only a modest increase in MAE relative to the full external set, whereas scaffold novelty was more damaging. Target-novel and double-cold subsets were the most difficult regression regimes, with markedly higher MAE, strongly negative *R*^2^, and wide confidence intervals. The uncertainty of these estimates reflects their smaller sample sizes, particularly for novel receptors, but the direction is clear: extrapolation across receptor identity is substantially more difficult than interpolation among receptors represented during training.

Data source contributed independently. BindingDB observations showed lower MAE than the ChEMBL 37 subset in the stratified analysis, demonstrating that “external” difficulty is not a simple function of chronological novelty. Assay provenance, endpoint composition, chemical coverage, and receptor coverage all contribute to the distribution shift.

Heterogeneity in the activity labels provides a plausible component of this behaviour. A common pActivity scale allows *K*_*i*_, *K*_*d*_, *IC*_50_, and *EC*_50_ measurements to be combined, but it does not make these quantities mechanistically identical or remove laboratory and assay-specific effects. Multi-task Bioassay Pre-training was motivated by precisely this problem: ChEMBL-scale affinity data contain inconsistent measurement types and experimental conditions that introduce systematic label noise [21]. The external gap observed here should therefore be interpreted as the joint effect of chemical novelty, target novelty, assay heterogeneity, and source/temporal drift rather than solely as failure to extrapolate molecular structure.

### 3.7 External Error Structure

No single measure of proximity to the training distribution explained the external errors. Maximum ligand similarity to the internal training compounds showed only a weak inverse association with absolute error (Spearman *ρ* = − 0.15; Fig. 4a). High structural similarity was therefore mildly favourable but far from sufficient: substantial errors occurred even for highly similar ligands, whereas many low-similarity compounds were predicted accurately. A simple Tanimoto threshold would consequently discard useful predictions while failing to identify all high-error cases.

**Fig. 4:**
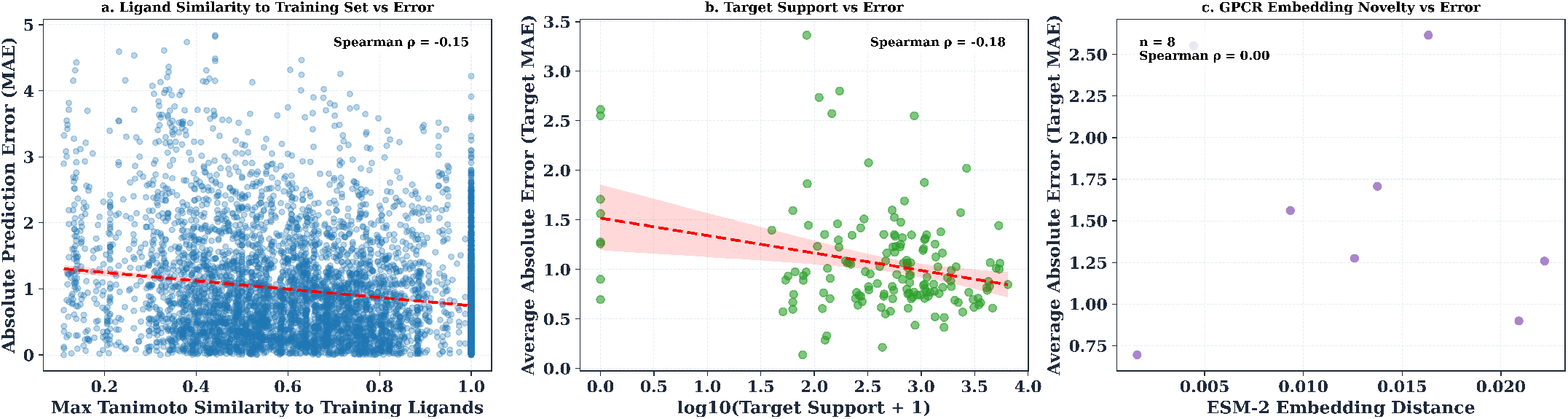
Relationship between external prediction error and training-domain proximity. (a) Absolute pActivity error versus maximum Morgan-fingerprint Tanimoto similarity to the training ligands. (b) Mean target-level absolute error versus GPCR training support. (c) Mean target-level error for target-novel GPCRs versus ESM-2 650M embedding distance from the nearest training receptor. Dashed lines indicate fitted trends; Spearman correlations are reported within each panel.

Target support behaved similarly. Mean target-level MAE decreased only weakly with increasing number of training observations (*ρ* = − 0.18; Fig. 4b). Additional ligand–target measurements improve coverage on average, but support count alone does not capture assay heterogeneity, local chemical diversity, or the pharmacological complexity of an individual receptor.

For the eight target-novel GPCRs, distance to the training receptors in ESM-2 embedding space showed no detectable rank association with error (*ρ* = 0.00; Fig. 4c). The sample is too small to infer the absence of a biological relationship, but the result does show that whole-sequence embedding distance was not an immediately useful applicability-domain criterion in this external set.

This last observation exposes a limitation of global mean-pooled receptor representations. A whole-protein embedding can distinguish receptors sufficiently well to improve activity and subtype-selectivity prediction, yet ligand recognition may depend on a small subset of binding-pocket or allosteric residues whose contribution is diluted in a global sequence vector. PSICHIC provides a useful contrast: its residue–ligand interaction fingerprints retain local sequence information and have been used to identify selectivity-relevant protein residues and ligand atoms [16].

The external results therefore favour an applicability framework based on predictive uncertainty and local ligand–receptor context, rather than one based solely on nearest-neighbour similarity. Conformal calibration, ensemble uncertainty, assay-aware training, or explicit residue-level ligand–protein interaction modelling are likely to provide more useful estimates of when a prediction lies outside the model’s reliable domain.

### 3.8 DRD2–DRD3 Selectivity

The strongest evidence that the receptor representation encodes pharmacologically meaningful subtype information came from the DRD2–DRD3 analysis. Despite the marked deterioration of absolute pActivity prediction on the broad external set, relative activity between these closely related dopamine receptors was recovered with high accuracy.

Across all 4,527 matched ligands, the cold-ligand ESM-2 650M ensemble achieved RMSE 0.447 (95% CI 0.431–0.461), MAE 0.323 (0.314–0.332), and *R*^2^ = 0.853 (0.842–0.863), with Spearman *ρ* = 0.916 and Pearson *r* = 0.924 (Table 6). Random-split performance was nearly identical, whereas scaffold-held-out compounds remained more difficult (**RMSE** = 0.501, *R*^2^ = 0.814). The same chemical-generalization hierarchy observed for absolute pActivity therefore persists for receptor selectivity, although the overall selectivity signal remains strong.

**Table 6:** DRD2–DRD3 selectivity performance of the ESM-2 650M CustomGNN. Regression statistics are seed-ensemble estimates with ligand-level bootstrap 95% confidence intervals. Classification evaluates strongly DRD3-selective compounds at Δ*p*Activity ≥ 1.

| Split | RMSE (95% CI) | MAE (95% CI) | R<sup>2</sup> (95% CI) | Spearman $\rho$ (95% CI) | Pearson $r$ (95% CI) | D3-selective ROC-AUC | D3-selective F1 |
| --- | --- | --- | --- | --- | --- | --- | --- |
| Random | 0.457 (0.441–0.472) | 0.332 (0.322–0.341) | 0.846 (0.834–0.856) | 0.912 (0.905–0.919) | 0.920 (0.914–0.926) | 0.964 (0.960–0.969) | **0.861 (0.852–0.871)** |
| Cold ligand | **0.447 (0.431–0.461)** | **0.323 (0.314–0.332)** | **0.853 (0.842–0.863)** | **0.916 (0.909–0.923)** | **0.924 (0.918–0.930)** | **0.966 (0.961–0.970)** | 0.856 (0.846–0.866) |
| Scaffold | 0.501 (0.484–0.519) | 0.366 (0.356–0.376) | 0.814 (0.802–0.827) | 0.895 (0.886–0.902) | 0.903 (0.896–0.910) | 0.956 (0.950–0.961) | 0.851 (0.840–0.861) |

The receptor ablation makes this result substantially stronger than the selectivity metrics alone. Without a GPCR embedding, RMSE increased to 1.552, *R*^2^ became −0.781, and rank correlations collapsed to approximately zero across the three principal splits. Classification of strongly DRD3-selective compounds likewise became indistinguishable from chance (ROC-AUC 0.487–0.499). The ligand representation can therefore reproduce aspects of the population-level chemistry, but it cannot infer whether an individual compound preferentially engages DRD2 or DRD3 in the absence of receptor information.

That distinction is visible in Fig. 5B. The observed fraction of compounds with Δ*p Activity* > 0 was 43.4%, closely reproduced by ESM-2 650M (43.1%), ProtT5 (41.4%), and even the no-ESM model (40.7%). Yet the ligand-only model fails at compound-level selectivity. Agreement between marginal distributions is therefore insufficient; receptor-aware pairwise prediction is what establishes subtype discrimination.

**Fig. 5:**
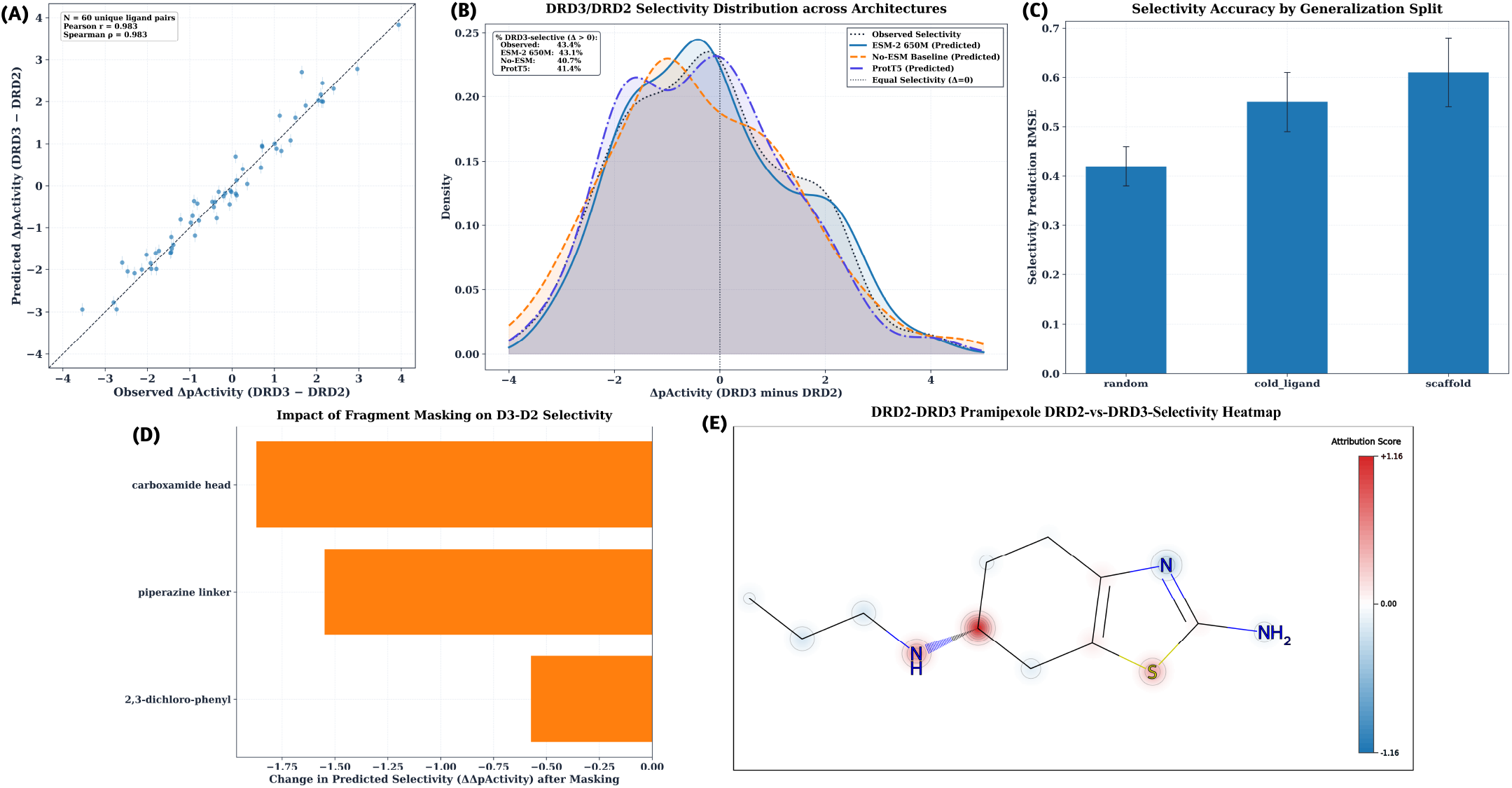
DRD2–DRD3 selectivity prediction and molecular interpretation. Selectivity was defined as Δ*p Activit y* = *p Activit y*_DRD3_ − *p Activit y*_DRD2_. (A) Observed versus predicted selectivity for a representative set of 60 held-out ligands using the three-seed ESM-2 650M ensemble; error bars indicate across-seed variability. (B) Observed and predicted selectivity distributions for ESM-2 650M, ProtT5, and the no-receptor-embedding model. (C) Selectivity RMSE across random, cold-ligand, and scaffold evaluation. (D) Change in predicted selectivity following masking of chemically defined Eticlopride fragments. (E) Differential atom attribution for Pramipexole, calculated as DRD3 attribution minus DRD2 attribution; red and blue indicate relative contributions toward DRD3- and DRD2-conditioned predictions, respectively.

The 60-ligand panel in Fig. 5A provides an illustrative compound-level view, with Pearson and Spearman correlations of 0.983. This subset is used for visualization only; the statistical estimates in Table 6 derive from the complete 4,527-ligand cohort. Split-resolved plots are provided in Fig. S5, and the representative-subset RMSEs in Fig. 5C reproduce the larger scaffold penalty observed in the full cohort.

D2/D3 selectivity is a particularly stringent receptor-aware test because the two receptors are highly homologous and subtype-selective ligand design remains non-trivial. Recent computational and medicinal-chemistry studies continue to identify secondary-pocket interactions and subtle ligand modifications as key determinants of D3 versus D2 preference [23], [24], [25], [26]. Won et al. recently reported dedicated D2/D3 QSAR models in which selectivity prediction outperformed individual receptor-affinity models and proposed cancellation of shared experimental noise after affinity subtraction as one contributing mechanism [23]. The present result extends that observation in an important respect. Selectivity was not learned by a receptor-pair-specific model; it emerged from two receptor-conditioned *p Activit y* predictions generated by the same multi-GPCR network. Shared errors can indeed cancel in the difference:

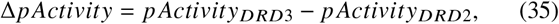

Δ*p Activit y* = *p Activit y*_*DRD*3_ − *p Activit y*_*DRD*2_, (35) but, error cancellation alone cannot explain the result because the no-receptor model remains at chance. The receptor embeddings therefore encode enough subtype-specific information for the shared network to alter its ligand predictions appropriately between DRD2 and DRD3.

### 3.9 Molecular Interpretation

The selectivity predictions were interrogated at two levels: chemically defined fragment perturbation and atom-level differential attribution.

Fragment masking produced highly asymmetric effects on predicted DRD3–DRD2 selectivity (Fig. 5D). Removing the carboxamide head altered Δ*p Activit y* by -1.872, and masking the piperazine linker by -1.549, compared with -0.573 for the 2,3-dichlorophenyl region. The selectivity prediction for this example is therefore not distributed uniformly across the molecular graph; the head group and linker carry substantially greater model sensitivity than the dichlorophenyl fragment.

This pattern is consistent with the medicinal chemistry of D2/D3-selective ligands, in which the orientation of an orthosteric pharmacophore and the chemistry of linkers or extended substituents can determine access to less conserved secondary binding regions. Zell et al. associated experimentally measured dopamine-receptor selectivity with secondary-pocket interactions, while Tian et al. showed that altering secondary binding fragments in fallypride-derived bitopic ligands changes D2/D3 affinity and selectivity [24], [25]. The correspondence is chemically plausible, but the fragment perturbation itself does not establish a physical receptor contact.

Atom-level differential attribution sharpened the same analysis. In Pramipexole, the predicted DRD3-versus-DRD2 contribution was concentrated in a small number of atoms rather than distributed uniformly across the molecule (Fig. 5E), with attribution scores spanning approximately −0.41 to +1.16. Additional maps for Eticlopride, Haloperidol, and SB-277011 show molecule-specific arrangements of positive and negative selectivity contributions (Fig. S6), arguing against a single universal atom-level selectivity motif.

The interpretation is necessarily model-specific. The receptor sequence is mean-pooled before ligand–protein fusion, and the model contains no explicit atom–residue interaction map. An attribution score therefore reports how a ligand feature influences the difference between predictions conditioned on DRD2 and DRD3 embeddings; it is not a binding energy, residue contact probability, or causal mechanism. Models such as PSICHIC, which explicitly learn residue–ligand interaction fingerprints, provide a route toward connecting predictive attribution more directly to receptor-side determinants [16].

The current interpretation nevertheless adds information beyond global feature importance. The model not only distinguishes DRD2 from DRD3 at cohort scale; perturbing defined ligand regions changes the predicted selectivity by more than one log unit in selected cases. These regions provide experimentally testable hypotheses for analogue design, while retaining the appropriate distinction between model sensitivity and mechanistic validation.

### 3.10 Implications and Limitations

The combined experiments define a clear hierarchy of what the model has learned. Ligand graphs provide a substantial advantage over fixed molecular features; receptor embeddings are essential for target-conditioned prediction; the custom message-passing formulation improves quantitative pActivity estimation beyond a matched GINE implementation; and chemically stringent partitioning reveals a scaffold-level extrapolation limit not apparent from random evaluation. These conclusions are supported across three independent seeds rather than by a single favourable training run.

The external results place equally important boundaries on those claims. Internal MAE of 0.51–0.64 increased to approximately 0.94–0.95 on ChEMBL 37/BindingDB, and no internal split strategy prevented this decline. Such a gap should not be treated as an inconvenient secondary result: it defines the actual applicability problem for prospective use. Recent zero-shot and cold-start DTI work likewise identifies transfer to unseen drugs, proteins, and domains as a central unresolved challenge [17], [22].

Part of this limitation is intrinsic to the supervision available in public bioactivity resources. *K*_*i*_, *K*_*d*_, *IC*_50_, and *EC*_50_ measurements capture related but non-equivalent pharmacological phenomena, and aggregation onto a common pActivity scale cannot remove assay-system or experimental-condition effects. The strong imbalance toward active observations further constrains learning of the negative interaction space. Assay-aware objectives and more extensive experimentally supported inactive data are therefore likely to be more valuable than simply increasing network capacity. The motivation for explicitly modelling assay heterogeneity in recent large-scale affinity learning supports this interpretation [21].

A second limitation lies in the receptor representation. Mean-pooled PLM embeddings offer an efficient structure-free route to modelling hundreds of GPCRs, and the DRD2–DRD3 results show that they preserve meaningful subtype information. They do not, however, encode ligand-dependent conformational states or explicit residue–atom interactions. Residue-level PLM representations, ligand–receptor cross-attention, structure-informed binding-site representations, or receptor conformational ensembles are logical next steps, particularly for the target-novel regime where whole-protein embedding distance was not predictive of error.

Finally, the DRD2–DRD3 analysis is a retrospective test of known activity differences, not prospective experimental validation. Its value is that it probes a biologically stringent form of receptor discrimination and provides interpretable hypotheses about which ligand regions influence that discrimination. Prospective use would require experimental testing of compounds prioritized by the model, particularly where predictions extend beyond the chemical or receptor domains represented in training.

The resulting picture is therefore deliberately more constrained than a claim of universal affinity prediction. The model performs strongly when chemical and receptor information can be learned jointly, retains substantial accuracy across unseen ligands and scaffolds, and encodes sufficient receptor-specific information to recover D2/D3 selectivity. Its principal remaining challenge is transfer beyond the bioactivity distribution on which that representation was learned. Addressing that challenge will require richer receptor–ligand interaction representations and explicit treatment of uncertainty and assay context rather than further optimization of internal benchmark performance alone.

## 4 Conclusion

We developed a multimodal graph-learning framework for GPCR ligand activity prediction that integrates explicit molecular graphs with pretrained receptor sequence representations and jointly models quantitative pActivity and binary activity. Across 271,739 curated ligand–GPCR pairs spanning 216 human GPCRs, the resulting CustomGNN consistently outperformed a protein-aware fixed-feature MLP and showed a reproducible regression advantage over a matched GINE reference architecture. Performance remained strong when exact ligands were withheld, whereas scaffold-disjoint evaluation revealed a larger loss of accuracy, identifying chemotype-level extrapolation as the principal internal generalization challenge.

Receptor information was central to this performance. Removing the protein-language-model representation substantially impaired both regression and classification, whereas differences among ESM-2 35M, ESM-2 650M, and ProtT5 were comparatively small. The selected ESM-2 650M model also recovered DRD2–DRD3 ligand selectivity with high accuracy, while the receptor-independent ablation approached chance-level subtype discrimination. This provides evidence that global sequence-derived representations retain pharmacologically relevant receptor-specific information even without explicit receptor structures.

Independent evaluation on ChEMBL 37 and BindingDB nevertheless exposed a substantial domain shift that was not captured by random, cold-ligand, or scaffold internal testing alone. This result defines an important boundary on prospective applicability and emphasizes that chemically stringent internal validation cannot substitute for independent temporal and source-level evaluation. Together, the findings establish protein-language-model-conditioned molecular graph learning as a scalable strategy for GPCR activity modelling, while identifying external-domain transfer, assay heterogeneity, and richer ligand–receptor interaction representations as the main priorities for further development.

